# Effects of local versus global competition on reproductive skew and sex differences in social dominance behaviour

**DOI:** 10.1101/2022.04.13.488121

**Authors:** Olof Leimar, Redouan Bshary

## Abstract

Social hierarchies are often found in group-living animals. The hierarchy position can influence reproductive success (RS), with a skew towards high-ranking individuals. The amount of aggression in social dominance varies greatly, both between species and between males and females within species. Using game theory we study this variation by taking into account the degree to which reproductive competition in a social group is mainly local to the group, emphasising within-group relative RS, or global to a larger population, emphasising an individual’s absolute RS. Our model is similar to recent approaches in that reinforcement learning is used as a behavioural mechanism allowing social-hierarchy formation. We test two hypotheses. The first is that local competition should favour the evolution of mating or foraging interference, and thus of reproductive skew. Second, decreases in reproductive output caused by an individual’s accumulated fighting damage, such as reduced parenting ability, will favour less intense aggression but should have little influence on reproductive skew. From individual-based simulations of the evolution of social dominance and interference, we find support for both hypotheses. We discuss to what extent our results can explain observed sex differences in reproductive skew and social dominance behaviour.

## Introduction

In group-living animals, positions in a social hierarchy are often established and maintained through pairwise aggressive interactions. The intensity of this aggression varies greatly, both between species and between males and females within species, with females typically showing less aggression than males [1, 2, 3]. There is also variation in the magnitude of reproductive skew caused by social dominance [4, 5, 3]. A traditional explanation for sex differences in reproductive skew and dominance behaviour is that there is greater scope for male than for female variation in reproductive success (RS), sometimes referred to as Bateman’s principle [1, 2, 3].

We analyse the evolution of reproductive skew and dominance behaviour by investigating the range from local to global competition in a metapopulation of local groups. The reasoning can apply either to males or to females. There is purely local competition if the group reproductive output is independent of reproductive skew, in which case a top-ranked individual in principle should have the capacity to make up a full group output. This individual would then have an incentive to monopolise reproduction in the group. For purely global competition, an individual’s RS should instead be measured against those in the larger population, so that there is little or no incentive to interfere with the reproductive output of other group members. The terms ‘soft selection’ and ‘hard selection’ are sometimes used for such a distinction between local and global competition [6]. For the evolution of reproductive skew and dominance behaviour, an important difference between local and global competition might then be that local competition favours mating or foraging interference (henceforth, interference). Interference can increase an individual’s relative RS in a group, whereas global competition only favours a high absolute RS. Concerning the evolution of sex differences in reproductive skew and dominance behaviour, evolution in males might be closer to local competition and in females to global competition, although with many intermediates between the extremes.

Apart from the scale of competition, various reproductive consequences of contest damage are likely to influence the evolution of dominance behaviour. Mortality from contest damage can decrease or eliminate reproduction and is one such effect. A reduced phenotypic quality that lowers parenting success could be a more widespread example, in particular in females [1, 2, 3]. Our aim is to elucidate the combined influence of the scale of competition (from local to global) and the costs of contest damage (in particular reduced parenting ability) on the evolution of dominance behaviour, interference, and reproductive skew. Our hypotheses are that interference is favoured by local competition and has a strong influence of the evolution of reproductive skew, and that other reproductive consequences of contest damage will influence the intensity of aggression, but will have a weaker effect on reproductive skew. As an alternative, we also consider situations where interference is very costly for dominants to perform and/or less effective in reducing subordinate reproduction, which should lead to less skew and less fighting. By investigating these hypotheses we aim to explain important aspects of sex-differences in reproductive skew and dominance behaviour. How local vs. global competition might influence the evolution of interference in social dominance has not previously been studied using game theory.

Our game-theory model is similar to previous approaches in using learning about differences in fighting ability as an evolving behavioural mechanism that can give rise to within-sex dominance hierarchies [7, 8, 9]. In addition, we introduce the strength of interference by dominants in subordinate reproduction as a trait that can evolve. The effect of variation in this trait spans from no interference to dominants nearly eliminating subordinate reproduction. For the case of no interference, we assume that a social hierarchy imposes a baseline level of reproductive skew, arising from such things as differences in the qualities of display arenas on a lek or of breeding sites, or possibly female preferences for males of different ranks. Interference can increase reproductive skew above this baseline.

In the following we outline the model elements and present results from individual-based evolutionary simulations. The genetically determined traits in the model are the components of a reinforcement learning mechanism, as used previously [8, 9], together with the strength of interference. We examine the evolution of these traits in one of the sexes, which could be either males or females. We then discuss to what extent our results provide a qualitative explanation of between-sex differences, and also if the factors we identify can throw light on within-sex species differences in reproductive skew and dominance behaviour. Finally, our analysis uses game theory to address the general question of why there is variation in the intensity of aggression, which was raised by Maynard Smith and Price [10] in their seminal contribution to game theory in biology, and we end by discussing sex differences in dominance behaviour from this perspective.

## The model

Our model here is an extension of previous models [8, 9], by adding variation in the degree of local competition and introducing interference as a trait. In previous models, competition was local, in the sense that dominance behaviour did not influence the total group reproductive output, but there was no interference, such that an individual’s RS was assumed to be directly determined by the dominance position it achieved. This means that in previous models, the amount of reproductive skew from social dominance was a model assumption and not a consequence of trait evolution. Here we introduce interference as a separate trait that influences an individual’s relative RS and study the co-evolution of this trait with other traits that determine the formation of a social hierarchy. Previously, we examined two types of costs of fighting damage, a decrease in the effective fighting ability and a risk of mortality from damage [9]. Here we add another cost of fighting, viz. a decreased parenting ability from fighting damage.

The elements of our model are outlined in Fig. 1. First a hierarchy is formed through aggressive interactions, then there is a risk of mortality from fighting damage, followed by reproduction and interference. Interference is a trait (*κ*) that measures how strongly an individual of a given dominance rank acts to reduce the reproduction of those of lower rank. Interference causes a proportional reduction in acquired resources (AR), i.e. the contested resources for reproduction acquired by a subordinate (Fig. 1b). Interference is costly to perform (Fig. 1b), and we assume that effects caused by different individuals interact multiplicatively. For both local and global competition, interference has the effect of increasing reproductive skew, but for global competition there is the additional effect of reducing the total group reproductive output (Fig. 1c). In addition to interference, we assume that accumulated contest damage can cause a proportional reduction in the individual’s parenting ability (Fig. 1d).

**Figure 1:**
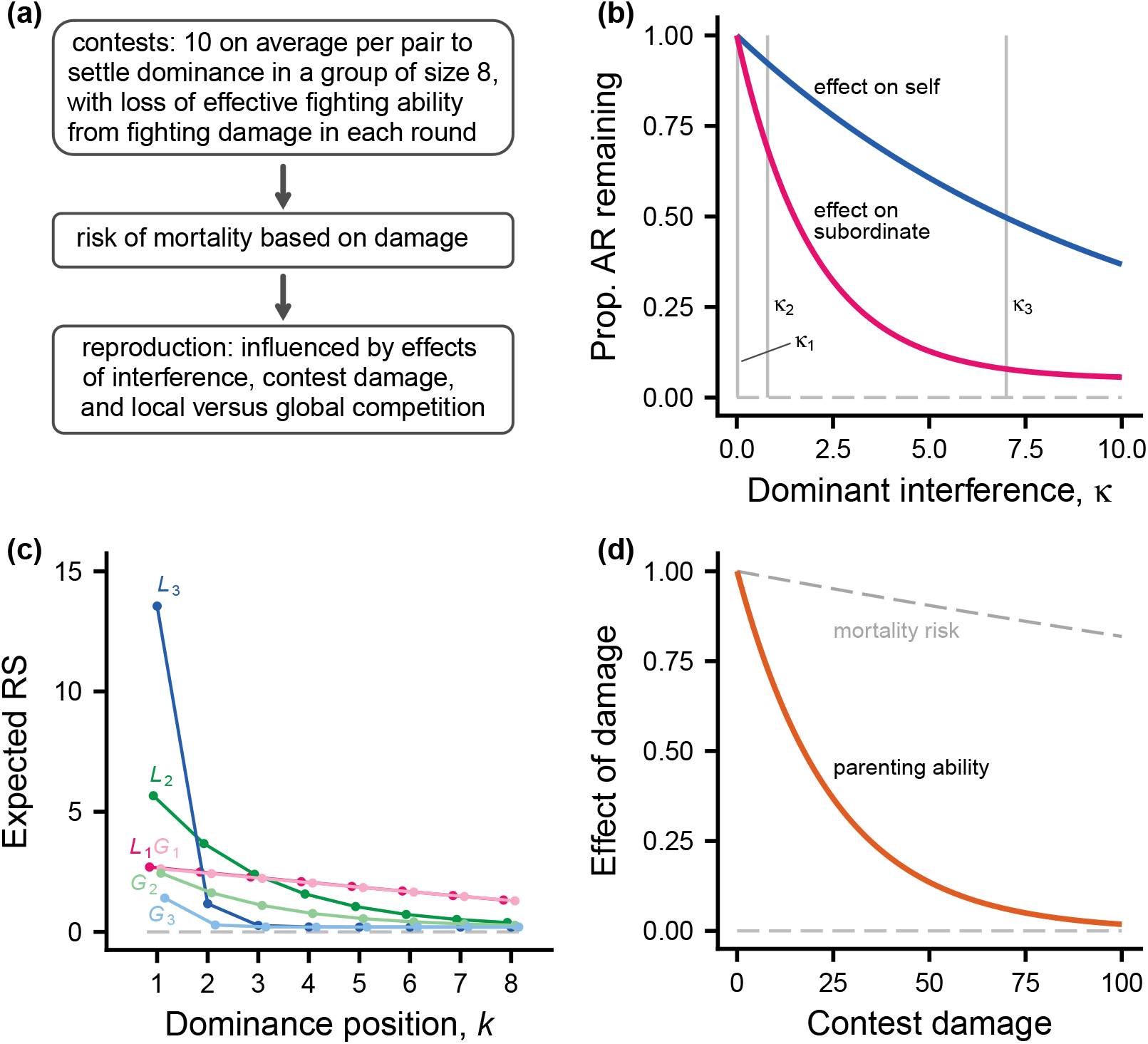
Illustration of model elements, with a one-season life cycle. The focus is on one of the sexes, either males or females. (a) Dominance-hierarchy formation comes first, followed by risk of mortality, which depends on contest damage, and reproduction. An individual’s reproductive success (RS) is the net result of different effects. (b) After hierarchy formation, dominant individuals can interfere with subordinates, reducing their acquired resources (AR), for instance matings for males or foraging opportunities for females. Interference strength is a genetically determined trait *κ*. The red curve shows the decrease in AR for a subordinate and the blue curve shows the decrease for a dominant from performing interference. The grey vertical lines indicate three values of *κ*. (c) Expected RS as a function of dominance position (*k* = 1 is top ranked) for local (*L*_1_, *L*_2_, *L*_3_) and global (*G*_1_, *G*_2_, *G*_3_) competition, corresponding to trait values *κ*_1_, *κ*_2_, *κ*_3_ from (b). The curves do not include effects of contest damage. Other local groups are assumed to have a total RS of 16 offspring. (d) The orange curve shows the influence of accumulated contest damage on parenting ability and the dashed curve shows the risk of mortality from contest damage.

To avoid the possibility that strong interference causes some individuals to be entirely without reproductive prospects, we assume that there is a small probability of ‘outside-option’ reproduction. The effect of this can be seen in Fig. 1c, where the curves labelled *L*_3_ and *G*_3_ come fairly close to, but do not reach zero for the bottom dominance positions (high values of *k*).

Many of the details of the model are the same as in previous models [8, 9], in particular the traits of the reinforcement-learning mechanism, but for completeness a full description is given in Supporting Information (SI), including a table of notation and definitions for the model (Table S1).

### Evolutionary simulations

Individuals are assumed to have genetically determined traits. The evolution of the traits is studied in individual-based simulations (Table 1). The traits for individual *i* include the strength of interference, *κ*_*i*_, and a number of traits of the reinforcement-learning mechanism. Of these, the degree of generalisation, *f*_*i*_, expresses how strongly an individual generalises learning about one opponent to other opponents, which is important for winner-loser effects. There are also the preference and value learning rates, *α*_*θi*_, *α*_*wi*_; the bystander learning rate, *β*_*i*_; the initial preference for the aggressive action A, *θ*_0*i*_; and the initial estimated value of a round, *w*_0*i*_. These are basic reinforcement-learning traits. Finally, the effect of observations on preference and value functions, *γ*_0*i*_, *g*_0*i*_, and the perceived reward from performing A, *v*_*i*_, are assumed to be genetically determined traits. See Table S1 and the SI text for further explanation.

**Table 1:** Trait values (mean ± SD over 100 simulations, each over 5000 generations) for 6 different cases of individual-based evolutionary simulations of social dominance interactions. The parameters that vary between cases are proportion local vs. global competition (*λ*) and the cost of contest damage (*d*).

| case | $\kappa_i$ | $f_i$ | $\alpha_{\theta i}$ | $\alpha_{wi}$ |
| --- | --- | --- | --- | --- |
| 1: $\lambda = 0.0, c_2 = 0.00$ | $0.020 \pm 0.005$ | $0.079 \pm 0.017$ | $52.2 \pm 6.7$ | $0.031 \pm 0.005$ |
| 2: $\lambda = 0.5, c_2 = 0.00$ | $0.799 \pm 0.160$ | $0.089 \pm 0.019$ | $54.5 \pm 7.6$ | $0.032 \pm 0.005$ |
| 3: $\lambda = 1.0, c_2 = 0.00$ | $6.759 \pm 0.330$ | $0.253 \pm 0.022$ | $22.5 \pm 6.1$ | $0.032 \pm 0.011$ |
| 4: $\lambda = 0.0, c_2 = 0.04$ | $0.020 \pm 0.004$ | $0.096 \pm 0.011$ | $134.8 \pm 17.1$ | $0.116 \pm 0.015$ |
| 5: $\lambda = 0.5, c_2 = 0.04$ | $0.603 \pm 0.147$ | $0.091 \pm 0.010$ | $124.9 \pm 14.3$ | $0.058 \pm 0.012$ |
| 6: $\lambda = 1.0, c_2 = 0.04$ | $6.510 \pm 0.423$ | $0.254 \pm 0.025$ | $52.1 \pm 10.5$ | $0.026 \pm 0.008$ |

| case | $\beta_i$ | $\theta_{0i}$ | $w_{0i}$ | $\gamma_{0i}$ | $g_{0i}$ | $v_i$ |
| --- | --- | --- | --- | --- | --- | --- |
| 1 | $1.10 \pm 0.18$ | $4.77 \pm 0.19$ | $-0.07 \pm 0.01$ | $1.20 \pm 0.27$ | $0.07 \pm 0.01$ | $0.82 \pm 0.01$ |
| 2 | $0.51 \pm 0.14$ | $4.90 \pm 0.15$ | $-0.02 \pm 0.01$ | $0.62 \pm 0.17$ | $0.04 \pm 0.01$ | $0.93 \pm 0.01$ |
| 3 | $0.18 \pm 0.14$ | $5.52 \pm 0.38$ | $0.01 \pm 0.03$ | $0.68 \pm 0.41$ | $0.05 \pm 0.02$ | $0.95 \pm 0.01$ |
| 4 | $2.33 \pm 0.18$ | $4.12 \pm 0.16$ | $-0.19 \pm 0.02$ | $2.50 \pm 0.22$ | $0.06 \pm 0.01$ | $0.57 \pm 0.02$ |
| 5 | $1.02 \pm 0.18$ | $5.14 \pm 0.22$ | $-0.08 \pm 0.02$ | $1.74 \pm 0.24$ | $0.06 \pm 0.01$ | $0.74 \pm 0.03$ |
| 6 | $0.18 \pm 0.11$ | $5.08 \pm 0.28$ | $-0.02 \pm 0.01$ | $0.61 \pm 0.29$ | $0.04 \pm 0.01$ | $0.94 \pm 0.01$ |

In evolutionary simulations, each trait is determined by an unlinked diploid locus with additive alleles. Alleles mutate with a probability of 0.002 per generation, with normally distributed mutational increments. The standard deviation of mutational increments for each trait was adjusted to ensure that simulations could locate evolutionary equilibria (as seen in Table 1, the evolved traits vary in scale, and mutational increments need to reflect the scale of trait variation).

A simulated population consisted of 500 groups of 8 individuals taking part in dominance interactions (either males or females), plus 8 individuals of the other sex, resulting in a total population size of *N* = 8000. Each interacting individual was assigned a quality *q*_*i*_, independently drawn from a normal distribution with mean zero and standard deviation *σ*_*q*_. As a simplification we assume that all offspring disperse globally over all groups, to form the adults of the next generation. For each case reported in Table 1, simulations were performed over intervals of 5000 generations, repeated at least 100 times, to estimate mean and standard deviation of traits at an evolutionary equilibrium.

### Standard parameter values

The following ‘standard values’ of parameters (Table S1) were used: proportion of local competition, *λ* = 0.0, *λ* = 0.5, or *λ* = 1.0; probability of outside-option reproduction, *Q* = 0.1; distribution of individual quality, *σ*_*q*_ = 0.50; damage cost parameters, *c*_0_ = 0.02, *c*_1_ = 0.0004, *c*_2_ = 0.00 or *c*_2_ = 0.04; interference parameters *b*_0_ = 0.1, *b*_1_ = 0.5, *φ* = 0.95; observations of relative quality, *a*_0_ = 0.707, *σ* = 0.50; perceived penalty variation, *σ*_*p*_ = 0.25. For these parameters, around 50% of the variation in the observations by individuals in a round is due to variation in relative fighting ability, *q*_*i*_ *− q*_*j*_.

## Results

The trait values that evolved in our individual-based simulations are shown in Table 1, for different degrees of local competition and absence vs. presence of a cost of decreased parenting ability from damage. The strength of interference (*κ*_*i*_) for the three different degrees of local competition (*λ* = 0.0, 0.5, 1.0) correspond approximately to the three values illustrated in Fig. 1b, with greater interference for higher degree of local competition (Table 1). In contrast, the evolved interference traits were similar for absence vs. presence of a cost of decreased parenting ability (*c*_2_ = 0.00 vs. *c*_2_ = 0.04, Table 1). These results are in accordance with our hypotheses.

Figure 2 shows different aspects of the outcome of dominance and interference interactions for the cases in Table 1. The degree of local competition (*λ*) had a strong effect on the distribution of RS over ranks (Fig. 2a) and thus on reproductive skew (Fig. 2c), with higher skew when competition is more local, whereas the absence vs. presence of a cost of decreased parenting ability only weakly influenced these measures (Fig. 2a, c). In contrast, both more local competition and the absence of a parenting-ability cost of damage lead to higher contest damage (Fig. 2b). These results support the hypotheses we set out to test. In addition, contest damage tended to be higher for lower-ranked individuals, in particular without a cost of decreased parenting ability (Fig. 2b).

**Figure 2:**
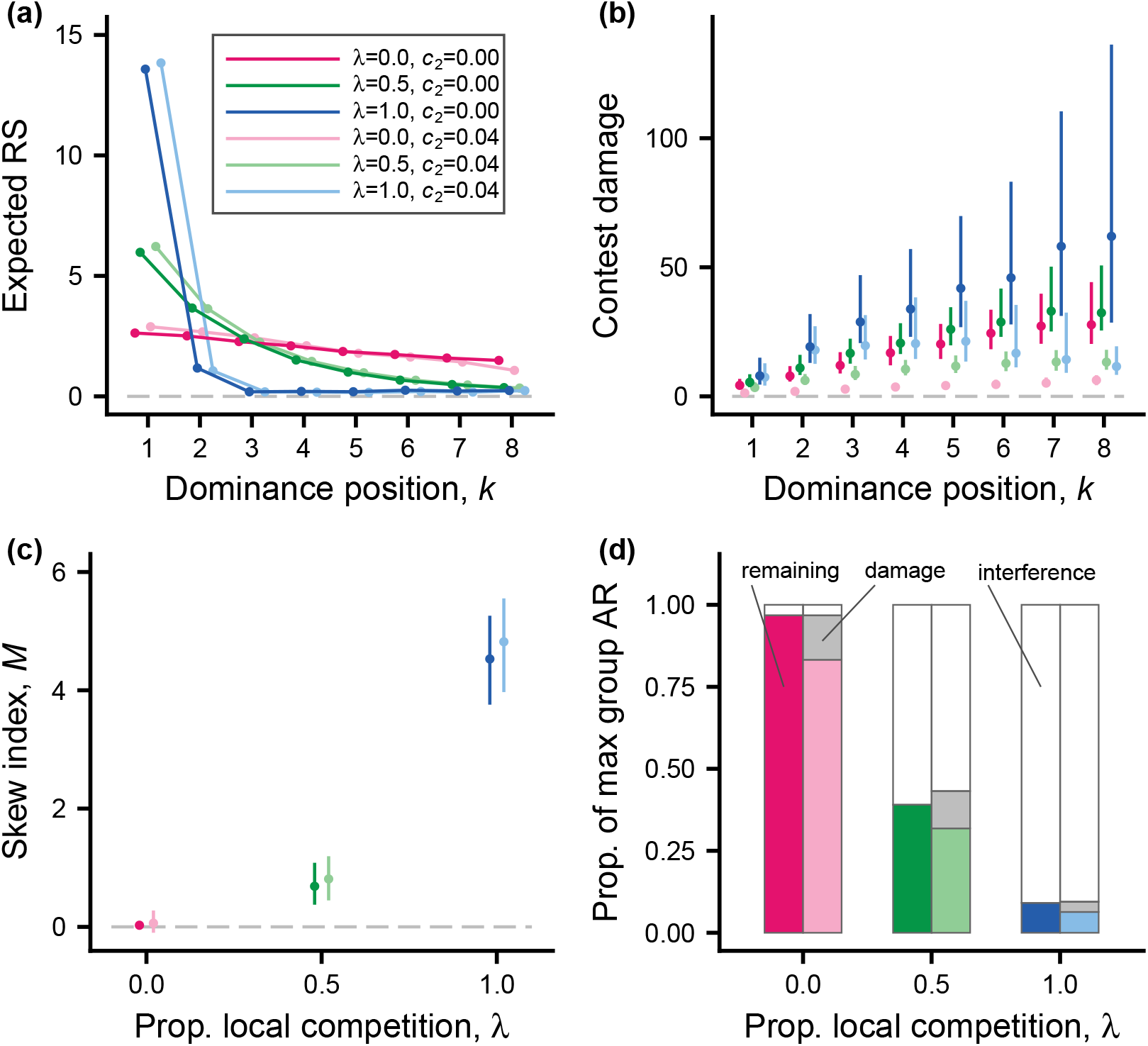
The outcome of individual-based simulations, showing effects of local vs. global competition and contest damage. (a) Expected RS as a function of dominance position for the 6 cases in Table 1 (colour coding given in legend). Red, green, and blue show global, intermediate, and local competition, respectively, with risk of mortality from Fig. 1d. The light-coloured curves in addition have reduced parenting ability from contest damage, as given in Fig. 1d. (b) Accumulated contest damage as a function of dominance position, for the 6 colour-coded cases in (a). Points and bars give median and 1st and 3rd quartiles for the distribution over simulated local groups. (c) Median and 1st and 3rd quartiles for the distribution over groups of the skew index *M*, as a function of the proportion of local competition (the 6 colour coded cases are shown). (d) Effects of interference and contest damage on the total group contested AR (summed over competing individuals), expressed as proportions of the maximal possible value. Unfilled parts indicate costs of interference, light grey parts costs of damage (reduced parenting ability), and colour-coded parts the remaining AR, respectively. Panels (a) and (b) include only surviving individuals; these have a dominance position. There were 500 groups in the simulated populations.

A comparison of the total group AR contributed by the competing sex for the different cases in Table 1 appear in Fig. 2d. In the cases with full local competition (*λ* = 1), interference strongly decreased the group AR. To interpret this, one can note that for full local competition, members of the competing sex (e.g., males) only contribute matings, but no additional resources to offspring. These instead come from the other sex (e.g., females). The sharp decrease in AR with full local competition thus only means that interference prevents most members of the competing sex from achieving contested matings. The interpretation of the cases with intermediate degree of local competition (*λ* = 0.5), and intermediate strength of interference, could instead be that a substantial part of the AR contributed by the competing sex is not subject to interference (e.g., nesting sites), but that interference can exclude individuals from other substantial parts (e.g., foraging areas). With global competition (*λ* = 0.0), there is very little interference and the only noticeable decrease in group AR comes from a small reduction in parenting ability from fighting damage (Fig. 1d).

The amount of fighting for the different ranks, and for local vs. global competition and absence/presence of a parenting-ability cost of damage is shown in Fig. 3, with examples of single groups in Fig. S1. There is much variation in the number of fighting rounds between individuals in different groups, but the tendency is that intermediate ranks fight the most. The tendency for the lowest ranks to fight less is an example of an ‘opt-out loser effect’, which we studied previously [9]. Part of the variation in fighting is that some pairs of individuals did not fight at all (Fig. S2). This was most common for global competition with a parenting-ability cost, and tended to occur when one of the individuals had low rank and the opponent had a higher rank (Fig. S2).

**Figure 3:**
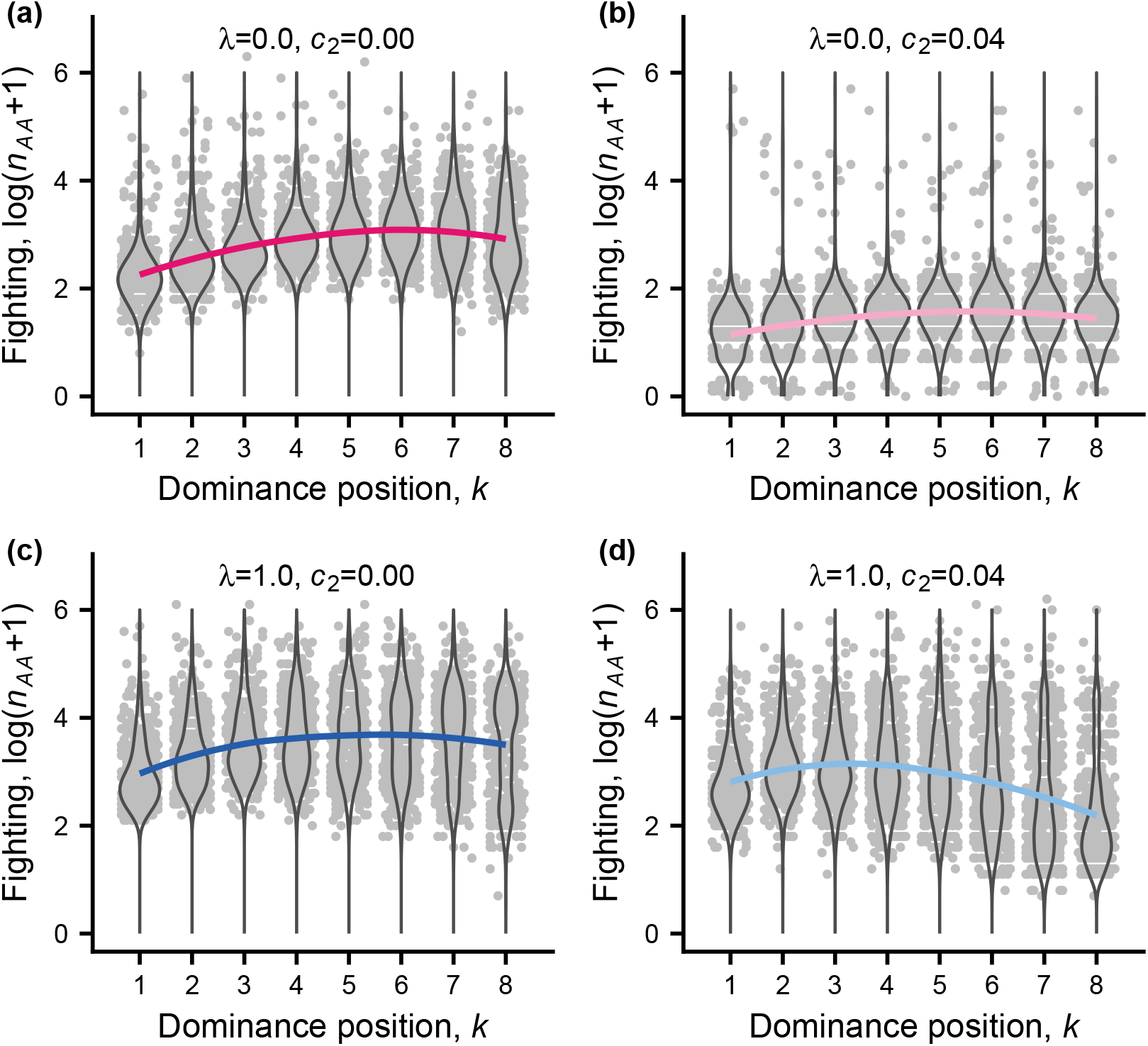
Simulated data (light grey points), density violins, and colour coded fitted curves of log-transformed total number of AA rounds (fighting rounds, loess fits) for cases with either no or full local competition. Panels (a), (b), (c), and (d) show cases 1, 3, 4, and 6 in Table 1, with colour coding as in Fig. 2. The x-values of the points are jittered for visibility.

The overall amount of fighting also varied substantially between the cases, being around 10 times lower for global competition with a parenting-ability cost than for local competition without parenting-ability cost (Fig. 3b vs. 3c).

We also investigated the evolutionary consequences of a substantially higher probability of outside-option reproduction (*Q* = 0.5, Table S2, Figs. S3, S4, S5). This did not strongly influence interference (see *κ*_*i*_ in Tables 1 and S2), but changed the learning traits in such a way that fights became shorter and less damaging (Fig. S3c and Figs. S4, S5). Because there was more outside-option reproduction, reproductive skew was reduced (Fig. S3a, c). With absence of a parenting ability cost, the lower ranks still tended to accumulate higher damage than the top ranks (Fig. S3b), but with full local competition and decreased parenting ability from damage the lowest ranks showed noticeably less fighting than the top ranks (Fig. S4c).

Finally, we examined the evolutionary consequences of a substantially more costly and less effective interference. As an example, for full local competition (*λ* = 1.0) and changing the interference parameters such that the lower curve in Fig. 1b gives the effect on self and the upper curve the effect on a subordinate, the outcome was that the interference traits evolved to near zero, leading to a baseline reproductive skew, and less fighting damage (Fig. S6).

## Discussion

Our evolutionary analysis showed that more intense local (within-group) competition favours stronger mating and/or foraging interference by dominants (Table 1), reducing the reproductive success of subordinates and increasing the reproductive skew (Fig. 2). We also found that costs in the form of reduced parenting ability from contest damage can sharply reduce fighting and the damage from fighting (Figs. 2, 3). These factors, separately or acting together, have effects that are large enough to potentially explain observed sex differences in social dominance behaviour. We further examined how fighting and damage varied between dominance positions, finding that individuals of intermediate or low ranks fought most and suffered the most contest damage (Figs. 2, 3).

The results on interference were achieved by introducing an evolving interference trait into the model. This is a new element compared to previous models of social-hierarchy formation that are similar to our current model in using reinforcement learning as a behavioural mechanism [8, 9]. Interference might correspond to different types of behaviours, ranging from dominant males attacking and chasing subordinate males to prevent them from mating, to dominant females excluding subordinate females from foraging areas. Among the examples are males of Alpine ibex, for which a dominance hierarchy is established before the start of the mating season [11, 12], and dominant, lactating olive baboon females excluding subordinate females from foraging through aggression [13]. The related idea of interference competition is much used in ecology, where it is applied to interactions both within and between species [14, 15] and can involve dominance interactions [16].

Interference, as used in our model, is related to punishment in animal societies, for which social dominance is a major example [17]. One idea is that punishment serves to deter cheating and promote cooperation, and another, contrasting idea is that it serves to change the relative RS in a group [18, 19]. In our model, interference plays the latter role, and can be costly by reducing the RS of the interfering individual. Interference resembles the concept of spite as used in theoretical studies on kin selection, with the conclusion that local competition favours spite [20]. Nevertheless, because we assume interacting individuals to be unrelated, interference in our model is not spite in the kin-selection sense [21].

In our model there are two contributions to reproductive skew: the ‘starting’ distribution of acquired resources (AR) over dominance ranks, which is the distribution for zero interference (*κ* = 0), and the change from this due to interference by dominants. Either of these contributions can vary between situations, giving rise to many possibilities. In addition, the cost and effectiveness of interference can vary, and if the cost becomes too high or effectiveness too low, interference will not evolve (see Fig. S6). This could, for instance, correspond to situations where the synchrony of receptivity of females in a group make it difficult of infeasible for high-ranking males to monopolise matings, as has been found in primate species [22].

To determine whether our model results explain observed sex-differences in social-dominance behaviour, one would need data on the scales of male and female competition in different species, as well as data on the influence of fighting damage on parenting ability, or similar effects of disturbance from fighting. Although these possible explanations have been put forward [1, 2], there seems to be a lack of quantitative estimates. Still, studies on female-female competition through interference in social groups support the general idea that interference is stronger when it can increase the RS of a dominant individual [23, 24, 13, 25].

Concerning the scale of competition, so-called female reproductive dominance is often assumed in life-history modelling (e.g., [26]). In species living in social groups, this would imply that male-male competition is mainly local, but there seem to be no studies directly investigating it. The concepts of hard and soft selection (the terminology is from [27]) are much used in studies of metapopulations [6] and correspond to global vs. local competition, but again there is little in the form of empirical estimates of these forms of population regulation. In the context of population management and conservation, data on culling, sterilization, or harvesting of either males or females are often studied (e.g., [28, 29, 30]), which in principle could allow estimates of the scale of competition, but up to now such data have not been used for this purpose.

One way to assess costs of dominance interactions on parenting ability is to examine genetic correlations between the corresponding traits, because such correlations could indicate an evolutionary trade-off. Evidence for such a trade off has been found in cows [31], with lower fertility and milk production in individuals more adapted for fighting. This complements the many observations consistent with decreased parenting ability from fighting [1, 2].

Hormonal manipulation is another way to examine such a trade-off, and this has been performed on cleaner fish in the wild [32]. Cleaner fish (*Labroides dimidiatus*) live on coral reefs. They forage by removing ectoparasites from other fish, so-called clients [33], and are organised into dominance hierarchies of females and a top-ranking male that has undergone sex reversal [34]. The study [32] found that testosterone-injected females increased their aggression towards subordinate females and spent less time interacting with clients, which supports our model assumptions. In general, cleaner fish, could be an example where female competition is relatively close to global, because they forage on non-monopolisable resources (clients) and reproduce through pelagic eggs.

There seems to be a lack of data on damage/fighting as a function of rank in social animals, but our result that high ranks suffer less damage has at least qualitative support from studies on health and longevity as a function of rank in social mammals [35]. Still, is has been argued [36] that improved health and longevity for high ranks typically applies to females, whereas high-ranking males males might suffer greater costs. So far there are no theoretical analyses explaining this potential sex difference. There are a number of previous models of reproductive skew, which have a focus on cooperative breeding among related individuals [37, 38, 39, 40, 41, 42], but these models could also apply to the situations we study here. So-called ‘tug-of-war’ models show some similarity to our approach, in allowing for different individual investments into conflict, but there are several notable differences. Previous reproductive skew models have not specifically dealt with either global vs. local competition, concrete mechanisms of hierarchy formation and interference, or allowed for more than two group members. Furthermore, our model does not make use of concepts like concessions, negotiations, or threats as possible explanations for sex-differences in social dominance behaviour [42], but instead uses reinforcement learning as a mechanism that allows hierarchy formation. Our model, as well as our previous model [9], makes relatively detailed assumptions and is therefore more complex than previous reproductive skew models, but it has the advantage of a somewhat closer match to field situations. This match might be further improved by incorporating more elements, such as age structure, relatedness between interacting group members, types of dispersal, and explicit modelling of outside-option reproduction.

Game theory in biology started 50 years ago as an attempt to explain why animal contests are often settled without serious injury [43, 10]. The originally proposed explanation, that threats of escalation and retaliation limit aggression at an evolutionary equilibrium [10, 44], has not stood the test of time, whereas the idea that assessment permits settlement with no or little fighting [45] is better supported. Sex differences in aggression and fighting in social animals might in fact be one of the main areas where the general question can be explored, adding to the interest in the problem. Our analysis here suggests that the life-history context, including such things as the scale of competition and reproductive opportunities outside of a given contest, is the main factor explaining how costly contests become. A combination of ambitious empirical work, including comparative studies, and theoretical modelling might throw further light on the issue.

## Acknowledgements

This work was supported by a grant (2018-03772) from the Swedish Research Council to OL and a grant (310030_192673/1) from the Swiss National Science Foundation to RB.

## Supporting information

### Additional tables

**Table S1:** Definitions and notation for the model.

| notation | definition or explanation |
| --- | --- |
| AR | acquired resources for reproduction, from mating or foraging |
| RS | reproductive success (number of offspring) |
| $\lambda$ | proportion local competition ($0 \leq \lambda \leq 1$) |
| $V(k)$ | baseline (no interference) AR going to rank $k$ ($k = 1$ top) |
| $Q$ | probability of rank-independent outside option RS |
| $M$ | reproductive skew index from Ross et al. (2020) |
| $q_i$ | quality (fighting ability) of individual $i$ |
| $\mu_q, \sigma_q$ | mean and SD of (normal) distribution of quality; $\mu_q = 0$ |
| $\hat{q}_{it}$ | damage-adjusted quality of individual $i$ |
| A, S | available actions: A is aggressive, S is submissive |
| $D_{it}$ | accumulated fighting damage for $i$ up to time $t$; each AA round between $i$ and $j$ increases damage by $e^{-(\hat{q}_{it}-\hat{q}_{jt})}$ |
| $c_0$ | parameter for loss of (adjusted) quality cost: $\hat{q}_{it} = q_i - c_0 D_{it}$ |
| $c_1$ | mortality cost parameter; Fig. 1d, survival is $e^{-c_1 D_{it}}$ |
| $c_2$ | parenting cost parameter; Fig. 1d, parenting ability is $e^{-c_2 D_{it}}$ |
| $\kappa_i$ | strength of interference for individual $i$ |
| $b_0$ | interference parameter; Fig. 1b, effect on self $e^{-b_0 \kappa_i}$ |
| $b_1, \phi$ | interference; Fig. 1b, effect on subordinate $1 - \phi + \phi e^{-b_1 \kappa_i}$ |
| $h_{iit} = f_i \theta_{iit}$ | generalised component of preference for action A at time $t$ |
| $f_i, \theta_{iit}$ | degree of generalisation and learned weight for individual $i$ |
| $h_{ijt} = (1 - f_i) \theta_{ijt}$ | opponent specific component of preference for A at time $t$ |
| $\theta_{ijt}$ | learned weight in opponent-specific component $h_{ijt}$ |
| $\theta_{0i}$ | starting value of $\theta_{iit}$ and $\theta_{ijt}$ |
| $\xi_{ijt}$ | observation by $i$, meeting $j$ at time $t$, of relative quality |
| $a_0, \epsilon_{ijt}$ | weight on $q_i, q_j$, and random error in observation $\xi_{ijt}$ |
| $\sigma$ | SD of (normally distributed) random error $\epsilon_{ijt}$ |
| $\gamma_{0i}$ | slope parameter for $i$ in preference component $\gamma_{0i} \xi_{ijt}$ |
| $p_{ijt}$ | probability to use action A by $i$ when meeting $j$ at time $t$ |
| $l_{ijt}$ | logit of $p_{ijt}$, referred to as the preference for the action A, defined as $l_{ijt} = h_{iit} + h_{ijt} + \gamma_{0i} \xi_{ijt}$ |
| $\hat{v}_{ijt}$ | estimated value (reward) by $i$ when meeting $j$ at time $t$ |
| $w_{iit}, w_{ijt}$ | generalised and opponent-specific learned weights in $\hat{v}_{ijt}$ |
| $w_{0i}$ | starting value of $w_{iit}$ and $w_{ijt}$ |
| $g_{0i}$ | slope parameter in $\hat{v}_{ijt} = f_i w_{iit} + (1 - f_i) w_{ijt} + g_{0i} \xi_{ijt}$ |
| $R_{ijt}$ | perceived reward by $i$ when meeting $j$ at time $t$ |
| $v_i$ | perceived reward by $i$ of performing the aggressive action A |
| $e_{ijt}, \sigma_p$ | random influence $e_{ijt}$ with SD $\sigma_p$ in penalty from AA round between $i$ and $j$, given by $\exp(-\hat{q}_{it} + \hat{q}_{jt} + e_{ijt})$ |
| $\alpha_{\theta i}, \alpha_{w i}$ | learning rates for updates by $i$ of weights in $l_{ijt}$ and $\hat{v}_{ijt}$ |
| $\beta_i$ | bystander learning rate, similar to $\alpha_{\theta i}$ |

**Table S2:** Trait values (mean SD ± over 100 simulations, each over 5000 generations) for 6 different cases of individual-based evolutionary simulations of social dominance interactions, with the probability of outside-option reproduction of *Q* = 0.5. The parameters that vary between cases are proportion local vs. global competition (*λ*) and the cost of contest damage (*d*).

| case | $\kappa_i$ | $f_i$ | $\alpha_{\theta i}$ | $\alpha_{wi}$ |
| --- | --- | --- | --- | --- |
| 1: $\lambda = 0.0, c_2 = 0.00$ | $0.031 \pm 0.008$ | $0.165 \pm 0.031$ | $12.7 \pm 1.5$ | $0.052 \pm 0.015$ |
| 2: $\lambda = 0.5, c_2 = 0.00$ | $0.928 \pm 0.113$ | $0.054 \pm 0.012$ | $76.5 \pm 6.4$ | $0.058 \pm 0.009$ |
| 3: $\lambda = 1.0, c_2 = 0.00$ | $6.766 \pm 0.333$ | $0.232 \pm 0.029$ | $54.4 \pm 8.7$ | $0.059 \pm 0.015$ |
| 4: $\lambda = 0.0, c_2 = 0.04$ | $0.031 \pm 0.006$ | $0.096 \pm 0.009$ | $262.3 \pm 13.6$ | $0.288 \pm 0.051$ |
| 5: $\lambda = 0.5, c_2 = 0.04$ | $0.831 \pm 0.118$ | $0.166 \pm 0.011$ | $135.1 \pm 10.4$ | $0.140 \pm 0.020$ |
| 6: $\lambda = 1.0, c_2 = 0.04$ | $6.570 \pm 0.229$ | $0.561 \pm 0.025$ | $29.5 \pm 3.4$ | $0.010 \pm 0.002$ |

| case | $\beta_i$ | $\theta_{0i}$ | $w_{0i}$ | $\gamma_{0i}$ | $g_{0i}$ | $v_i$ |
| --- | --- | --- | --- | --- | --- | --- |
| 1 | $2.18 \pm 0.34$ | $2.55 \pm 0.31$ | $-0.05 \pm 0.03$ | $1.18 \pm 0.26$ | $0.10 \pm 0.02$ | $0.78 \pm 0.02$ |
| 2 | $0.38 \pm 0.12$ | $4.31 \pm 0.13$ | $-0.04 \pm 0.01$ | $1.06 \pm 0.18$ | $0.06 \pm 0.01$ | $0.85 \pm 0.01$ |
| 3 | $0.16 \pm 0.07$ | $4.15 \pm 0.18$ | $-0.04 \pm 0.02$ | $0.62 \pm 0.22$ | $0.06 \pm 0.02$ | $0.94 \pm 0.02$ |
| 4 | $3.59 \pm 0.28$ | $1.52 \pm 0.15$ | $-0.39 \pm 0.04$ | $3.21 \pm 0.17$ | $0.06 \pm 0.01$ | $0.37 \pm 0.04$ |
| 5 | $1.96 \pm 0.21$ | $3.71 \pm 0.17$ | $-0.19 \pm 0.02$ | $2.42 \pm 0.17$ | $0.08 \pm 0.02$ | $0.62 \pm 0.02$ |
| 6 | $1.65 \pm 0.27$ | $2.22 \pm 0.22$ | $-0.08 \pm 0.02$ | $1.08 \pm 0.14$ | $0.07 \pm 0.01$ | $0.71 \pm 0.02$ |

### Additional figures

**Figure S1:**
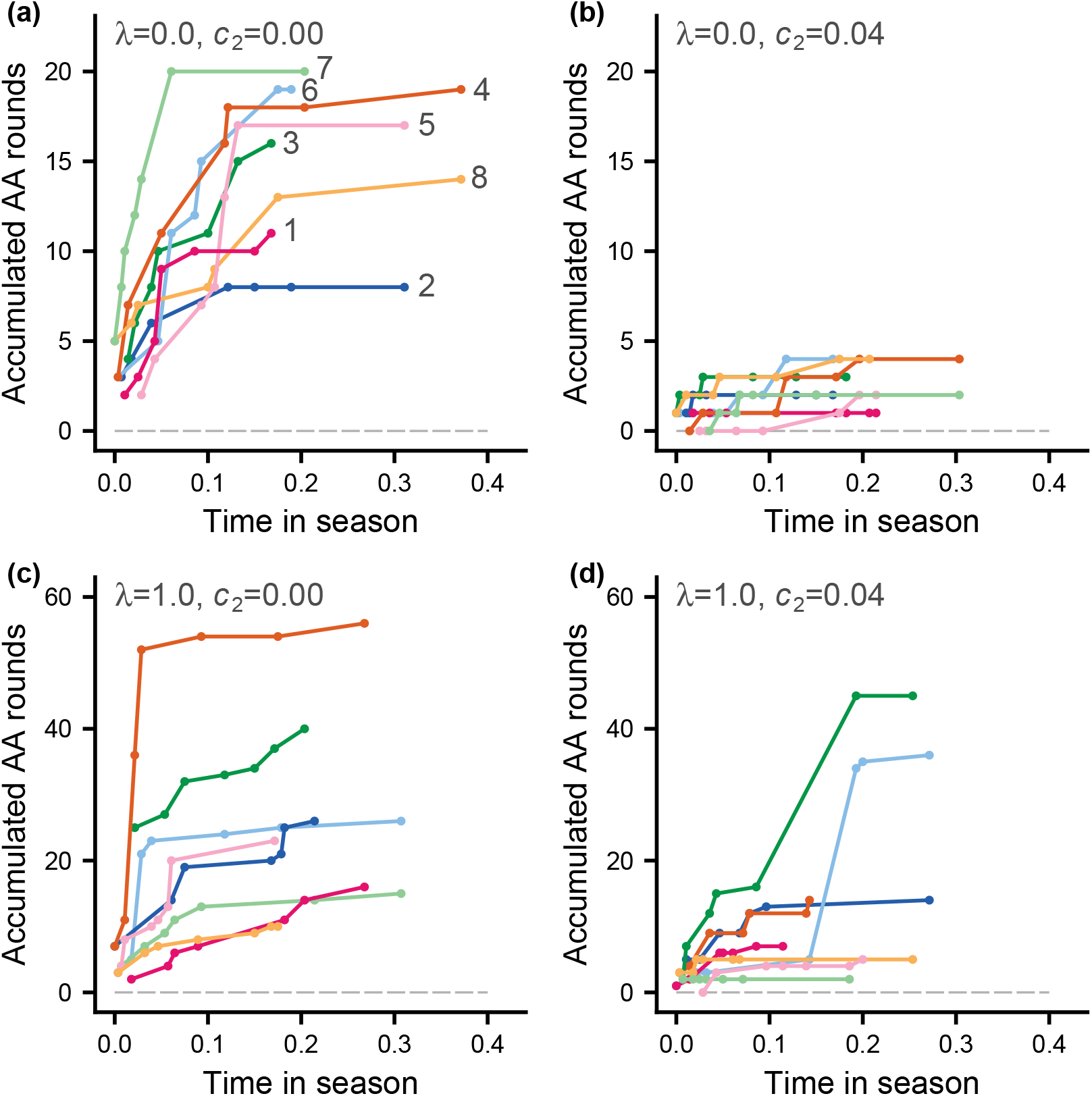
Accumulated number of AA rounds (fighting rounds) for individuals of different ranks as a function of time in the season. Each panel shows one group from the corresponding cases in Table 1 that are shown in the panels in Fig. 3. The curves are colour coded according to an individual’s rank, with labels in panel (a). Time on the x-axes has been defined such that 1.0 corresponds to completion of all contest. Note that the scales differ between the y-axes.

**Figure S2:**
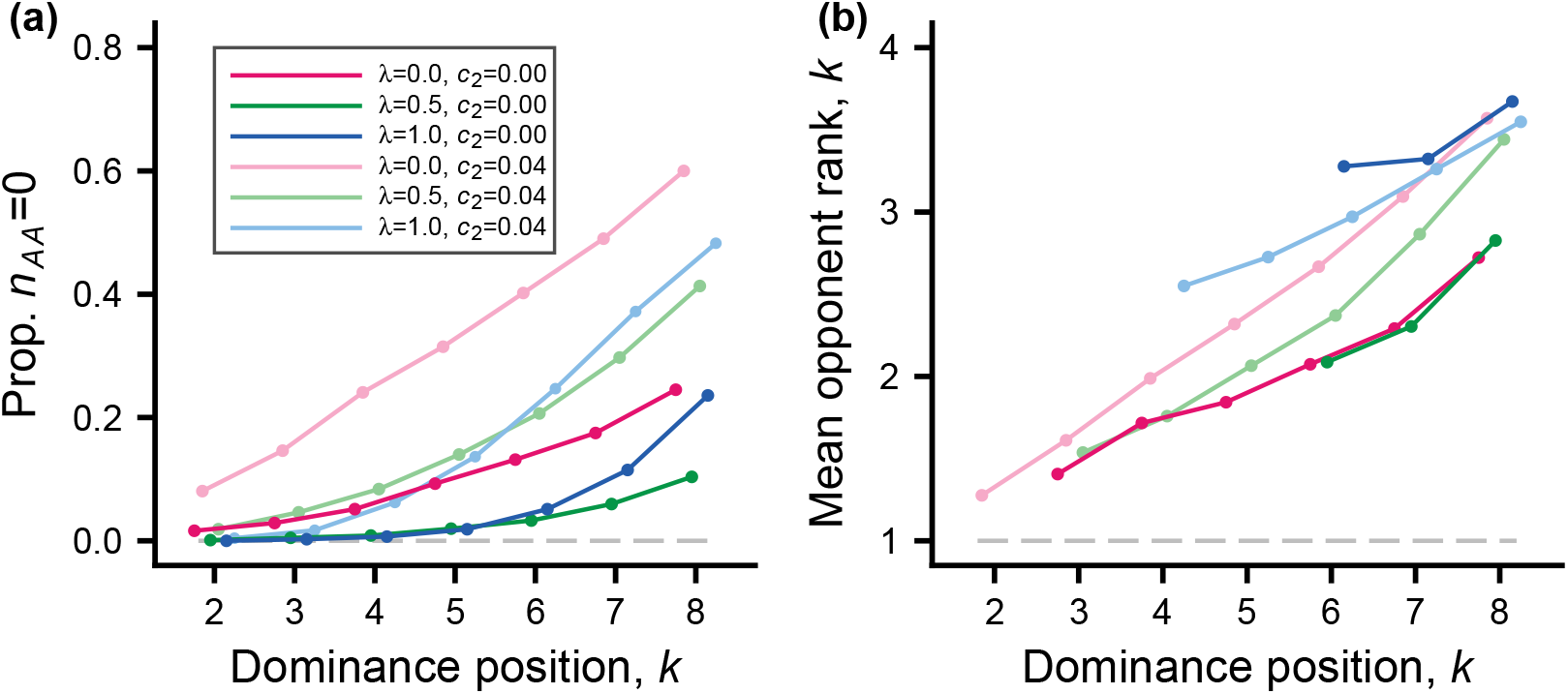
Illustration of contests where there is no fighting (number of AA rounds is zero), because one individual is submissive throughout the contest, for the cases in Table 1. Panel (a) shows the proportion of all contests for individuals of a certain rank where the individual avoided fighting against an opponent. Only data for rank positions 2 to 8 are shown; top-ranked-to-be individuals were rarely submissive throughout a contest. Panel (b) shows the mean rank of this opponent. Only data for rank positions in (a) with a proportion above 0.025 are shown in panel (b). Colour coding and underlying simulation data are the same as in Fig. 2.

**Figure S3:**
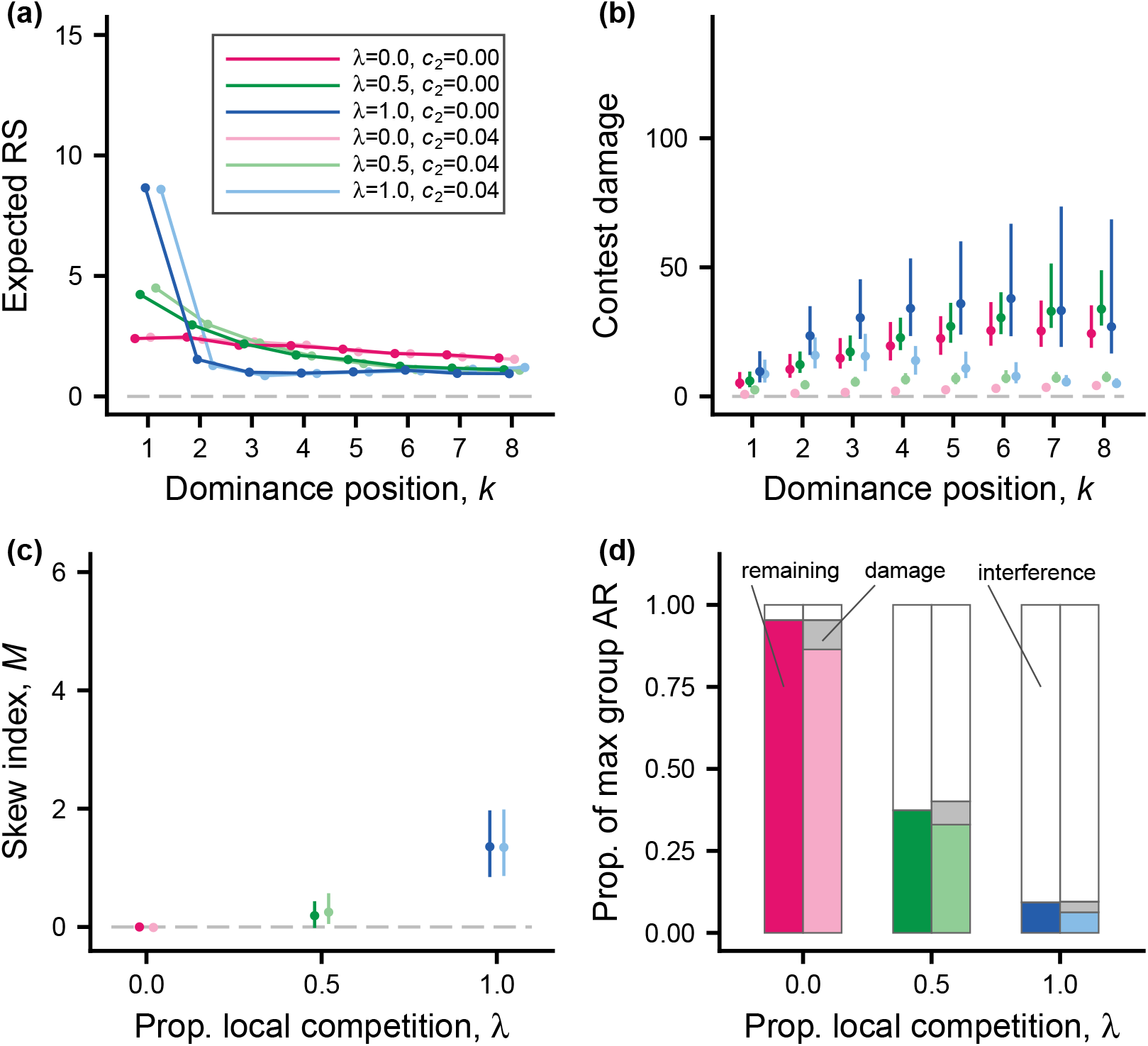
The outcome of individual-based simulations, showing effects of local vs. global competition and contest damage. The probability of outside-option reproduction is *Q* = 0.5. The figure can be compared with Fig. 2, where *Q* = 0.1. (a) Expected RS as a function of dominance position for the 6 cases in Table S2 (colour coding given in legend). Red, green, and blue show global, intermediate, and local competition, respectively, with risk of mortality from Fig. 1d. The light-coloured curves in addition have reduced AR from contest damage, as given in Fig. 1d. (b) Accumulated contest damage as a function of dominance position, for the 6 colour-coded cases in (a). Points and bars give median and 1st and 3rd quartiles for the distribution over simulated local groups. (c) Median and 1st and 3rd quartiles for the distribution over groups of the skew index *M*, as a function of the proportion of local competition (the 6 colour coded cases are shown). (d) Effects of interference and contest damage on the total group contested AR (summed over competing individuals), expressed as proportions of the maximal possible value. Unfilled parts indicate costs of interference, light grey parts costs of damage (reduced parenting ability), and colour-coded parts the remaining AR, respectively. Panels (a) and (b) include only surviving individuals; these have a dominance position. There were 500 groups in the simulated populations.

**Figure S4:**
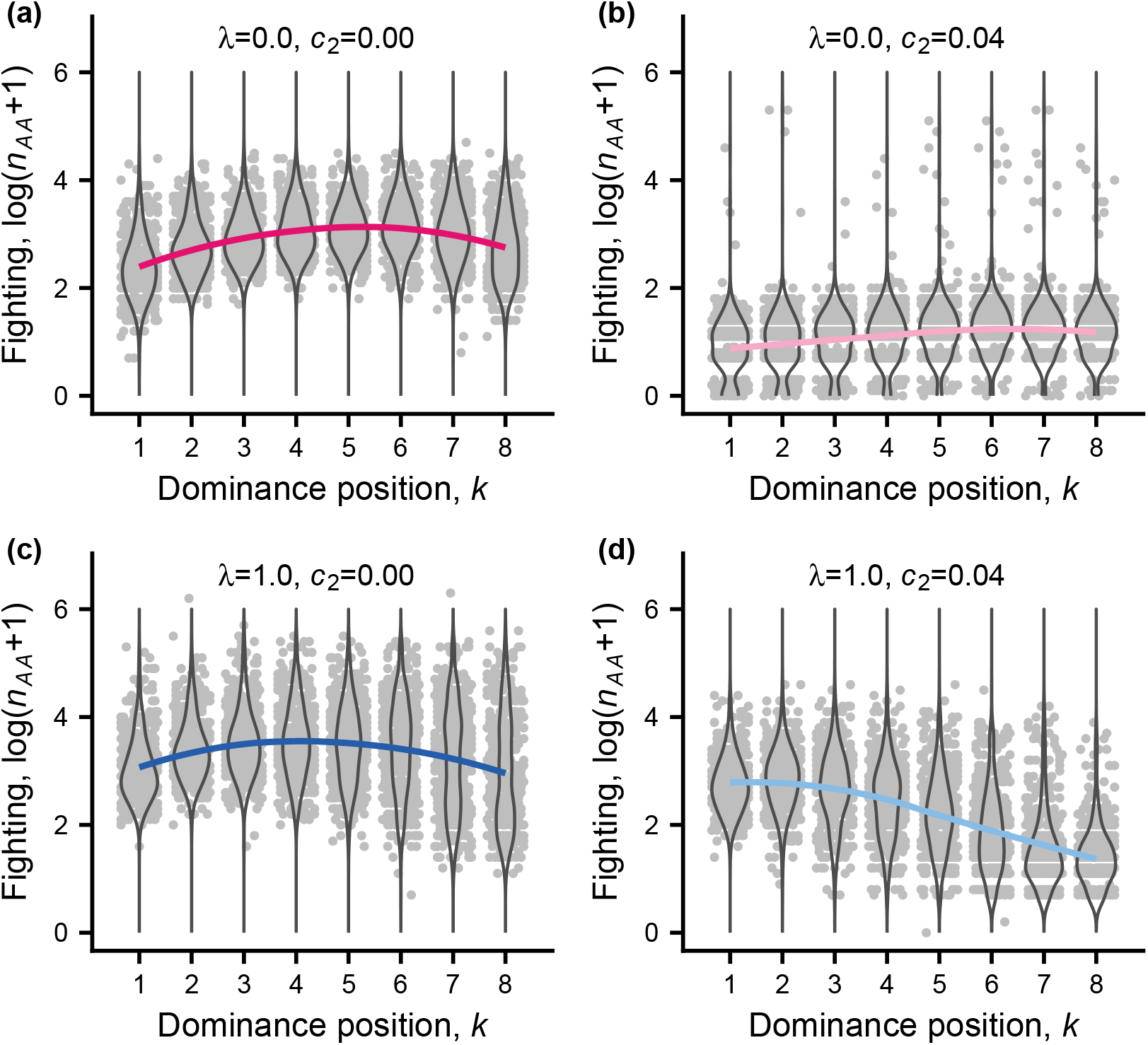
Simulated data (light grey points), density violins, and colour coded fitted curves of log-transformed total number of AA rounds (fighting rounds, loess fits) for cases with either no or full local competition. Panels (a), (b), (c), and (d) show cases 1, 3, 4, and 6 in Table S2 (*Q* = 0.5), with colour coding as in Fig. S3. The x-values of the points are jittered for visibility. The figure can be compared with Fig. 3, where *Q* = 0.1.

**Figure S5:**
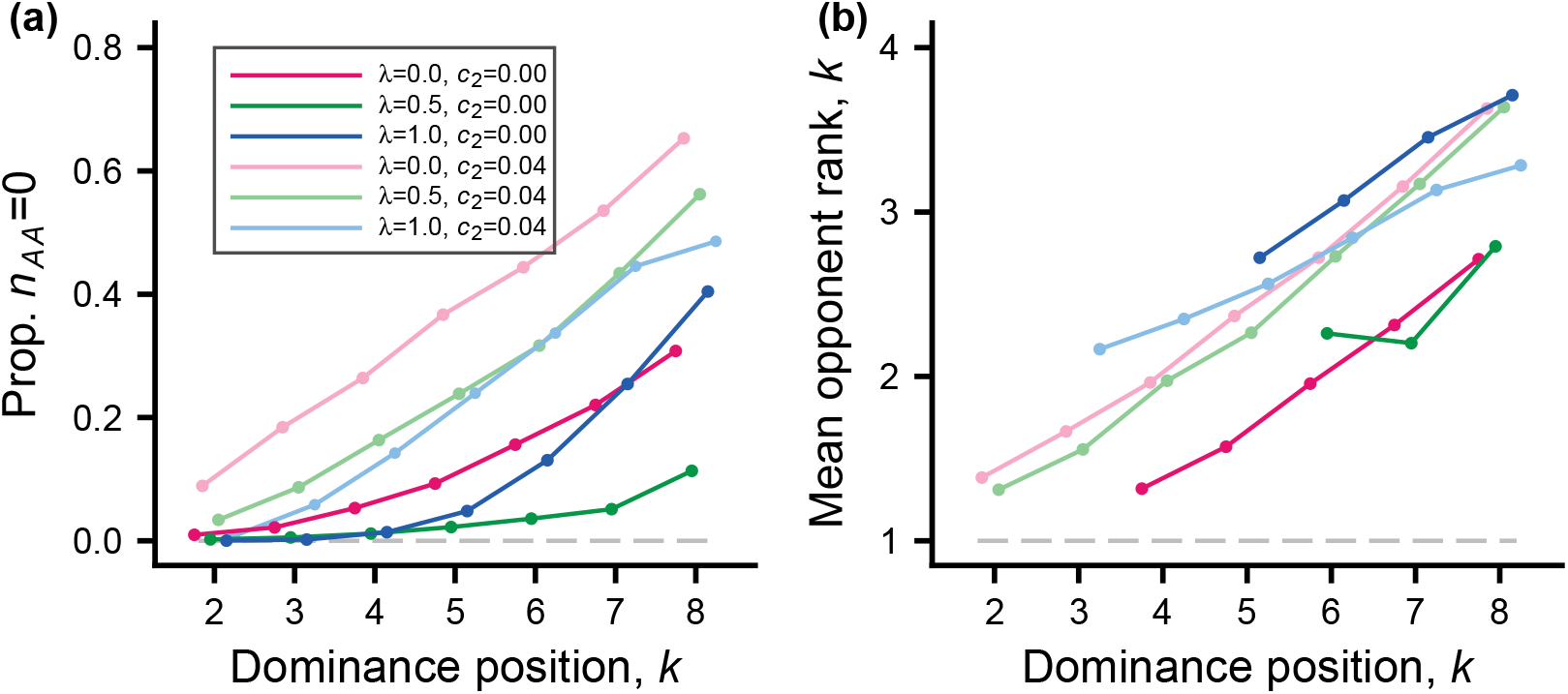
Illustration of contests where there is no fighting (number of AA rounds is zero), because one individual is submissive throughout the contest, for the cases in Table S2. Panel (a) shows the proportion of all contests for individuals of a certain rank where the individual avoided fighting against an opponent. Only data for rank positions 2 to 8 are shown; top-ranked-to-be individuals were rarely submissive throughout a contest. Panel (b) shows the mean rank of this opponent. Only data for rank positions in (a) with a proportion above 0.025 are shown in panel (b). Colour coding and underlying simulation data are the same as in Fig. S3.

**Figure S6:**
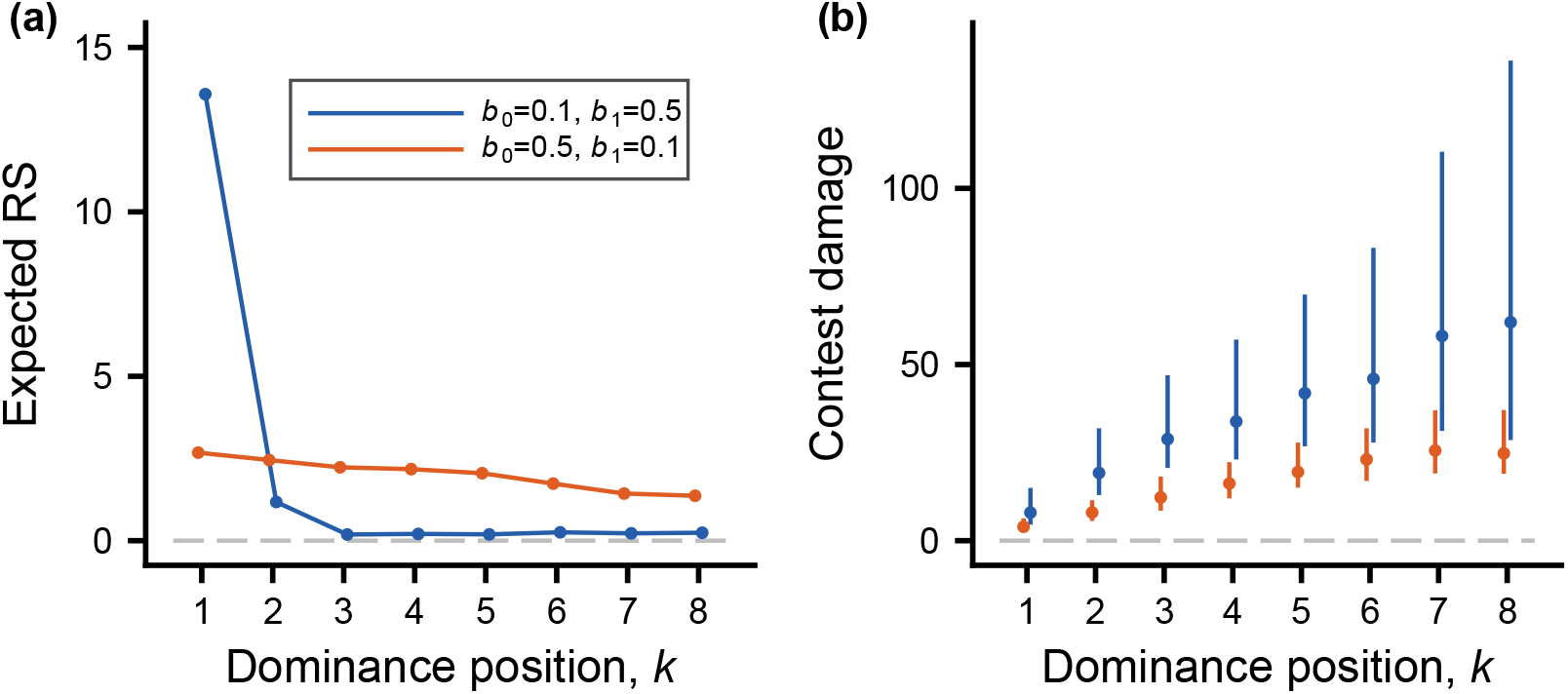
The outcome of individual-based simulations, illustrating the evolutionary consequences of interference being more costly and less effective. Two cases are illustrated. One is the same as the case with *λ* = 1.0 (i.e., local competition) and *c*_2_ = 0.00 in Fig. 2 (dark blue colour); this case has interference parameters *b*_0_ = 0.1, *b*_1_ = 0.5 (see Table S1), which correspond to the curves in Fig. 1b. In the second case (dark orange colour), the interference parameters are changed to *b*_0_ = 0.5, *b*_1_ = 0.1, making interference more costly and less effective and leading to interference traits *κ*_*i*_ evolving to approximately zero. (a) Expected RS as a function of dominance position. (b) Accumulated contest damage as a function of dominance position. Points and bars give median and 1st and 3rd quartiles for the distribution over simulated local groups. The figure includes only surviving individuals; these have a dominance position. There were 500 groups in the simulated populations. The figure can be compared with panels (a) and (b) in Fig. 2.

## Model details

Brief descriptions of model elements appear in Table S1. Several elements of the model are similar to those in previous models [8, 9]. These are the observations in a round, including assumptions about individual recognition, the actions A and S, the action preferences and estimated values, the implementation of action exploration, and the learning updates, including bystander learning.

### Life-history, competition and reproductive season

There is an annual life cycle with a single reproductive season (Fig. 1a). Dominance interactions occur in groups with *g*_*s*_ members of the competing sex (*g*_*s*_ = 8 for the individual-based simulations in Table 1). The season starts with a sequence of contests. Each contest is between a randomly selected pair of group members and there are 5*g*_*s*_(*g*_*s*_ *−* 1) contests, i.e., on average 10 contests per pair, but as soon as dominance has been established for a pair, there is no further fighting in that pair. An individual’s survival from the contests to reproduction depends on its accumulated fighting damage. As a result of the contests, a dominance hierarchy is formed, and surviving group members obtain contested reproductive success (RS) according to their acquired resources (AR), in a way that is influenced by the proportion *λ* of local competition. The interpretation of *λ* is that for pure local competition (*λ* = 1) each group has the same expected total contested RS, whereas for pure global competition (*λ* = 0) the entire population of parents produce sufficient offspring for the next generation and each competing parent’s expected contested RS is proportional to its AR. For intermediate competition (0 *< λ <* 1), the expected total contested RS of a group is proportional to *λ*AR_tot_*/G* + (1 *− λ*)AR_loc_, where AR_tot_ is the total AR in the population, *G* is the number of groups, and AR_loc_ is the total AR of the group in question. Each competing individual in the group contributes to this group total in proportion to its AR. Individuals of the non-competing sex have equal chances of contributing to reproduction.

In addition to contested RS, the model implements an ‘outside option’ of non-contested RS. With a probability *Q* (with *Q* = 0.1 and *Q* = 0.5 for the simulations in Table 1 and Table S1 respectively) the parents of an offspring are randomly selected among all the living adults, without dependence on AR, and with probability 1 *− Q* the parents are selected as described above for contested RS. A parent of the competing sex needs to survive to contribute to outside-option reproduction, and for cases with non-zero parenting cost of damage (*c*_2_ *>* 0) the outside-option contribution is proportional to the parenting ability. Thus, the effects depicted in Fig. 1d are taken into account in outside-option RS. The reason for having outside-option RS is to avoid the extreme case of certain individuals having no reproductive future, which could otherwise happen given our assumption of an annual life-cycle and the possibility of strong interference by dominants in subordinate reproduction. For instance, for the cases labelled *L*_3_ and *G*_3_ in Fig. 1c, the expected RS is close to 0.2 (i.e., 2*Q*) for the bottom rank (*k* = 8), whereas it would have been close to zero without an outside option (see also Fig. S3.

We assume that offspring disperse randomly over the population. More details on hierarchy formation and reproduction appear below.

### Observations and actions

The model simplifies a round of interaction into two stages. In the first stage, interacting individuals make an observation. Thus, individuals observe some aspect *ξ* of relative fighting ability and also observe the opponent’s identity. The observation by an individual is statistically related to the difference in fighting ability between itself and the opponent, *q*_*i*_ *− q*_*j*_. For the interaction between individuals *i* and *j* at time *t*, the observation is

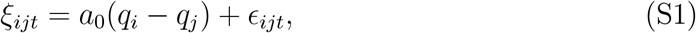

where *a*_0_ *>* 0 and *E*_*ijt*_ is an error of observation, assumed to be normal with mean zero and SD *σ*. Note here that the observations *ξ*_*ijt*_ refer to the original fighting abilities *q*_*i*_ and *q*_*j*_, and not the effective fighting abilities (see below).

By adjusting the parameters *σ*_*q*_, which is the SD of the distribution of *q*_*i*_, and *a*_0_ and *σ* from equation (S1), one can make the information about relative quality more or less accurate. The observation (*ξ*_*ij*_, *j*) is followed by a second stage, where individual *i* chooses an action, and similarly for individual *j*. The model simplifies to only two actions, A and S, corresponding to aggressive and submissive behaviour.

### Action preferences and estimated values

For an individual *i* interacting with *j* at time *t, l*_*ijt*_ denotes the preference for A. The probability that *i* uses A is then

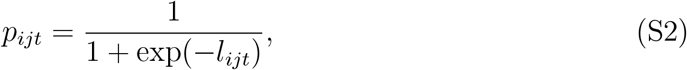

so that the preference *l*_*ijt*_ is the logit of the probability of using A. The model uses a linear (intercept and slope) representation of the effect of *ξ*_*ijt*_ on the preference, and expresses *l*_*ijt*_ as the sum of three components

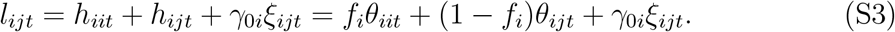

Here *h*_*iit*_ = *f*_*i*_*θ*_*iit*_ is a contribution from generalisation of learning from all interactions, *h*_*ijt*_ = (1 *− f*_*i*_)*θ*_*ijt*_ is a contribution specifically from learning from interactions with a particular opponent *j*, and *γ*_0*i*_*ξ*_*ijt*_ is a contribution from the current observation of relative fighting ability. Note that for *f*_*i*_ = 0 the learning about each opponent is a separate thing, with no generalisation between opponents, and for *f*_*i*_ = 1 the intercept component of the action preference is the same for all opponents, so that effectively there is no individual recognition (although the observations *ξ*_*ijt*_ could still differ between opponents). One can similarly write the estimated value 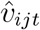 of aninteraction as a sum of three components:

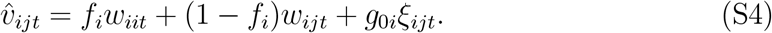

The actor-critic method updates *θ*_*iit*_, *θ*_*ijt*_, *w*_*iit*_, and *w*_*ijt*_ in these expressions based on perceived rewards, whereas *f*_*i*_, *γ*_0*i*_, and *g*_0*i*_ are genetically determined.

### Exploration in learning

For learning to be efficient over longer time spans there must be exploration (variation in actions), in order to discover beneficial actions. Learning algorithms, including the actor-critic method, might not provide sufficient exploration [46], because learning tends to respond to short-term rewards. In the model, exploration is implemented as follows: if the probability in equation (S2) is less than 0.01 or greater than 0.99, the actual choice probability is assumed to stay within these limits, i.e. is 0.01 or 0.99, respectively. In principle the degree of exploration could be genetically determined and evolve to an optimum value, but for simplicity this is not implemented in the model.

### Fighting damage and effective fighting ability

A group member *i* accumulates damage *D*_*it*_ from fighting. *D*_*it*_ refers to accumulated damage up to (but not including) round *t*. As a consequence of the damage, the individual’s effective fighting ability is reduced from the original *q*_*i*_ to

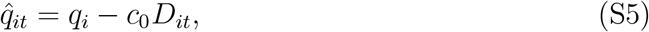

where *c*_0_ is a parameter. Following an AA round between *i* and *j*, there is an increment to *D*_*it*_:

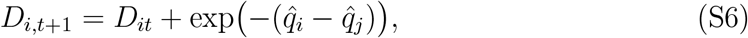

and similarly for *j*. The effective fighting abilities also determine the perceived costs (see below), and in this way they influencing the learning.

### Perceived rewards

An SS interaction is assumed to have zero rewards, *R*_*ijt*_ = *R*_*jit*_ = 0. For an AS interaction, the aggressive individual *i* perceives a reward *R*_*ijt*_ = *v*_*i*_, which is genetically determined and can evolve. The perceived reward for the submissive individual *j* is zero, *R*_*jit*_ = 0, and vice versa for SA interactions. If both individuals use A, some form of costly interaction or fight occurs, with perceived costs (negative rewards or penalties) that are influenced by the effective fighting abilities of the two individuals. The perceived rewards of an AA interaction are assumed to be

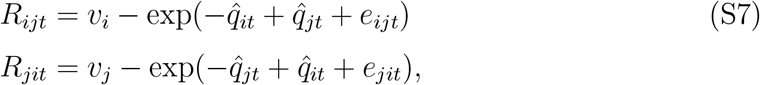

where *e*_*ijt*_ is a normally distributed random influence on the perceived penalty, with mean zero and standard deviation *σ*_*p*_, and similarly for *e*_*jit*_.

### Learning updates

In actor-critic learning, an individual updates its learning parameters based on the prediction error (TD error)

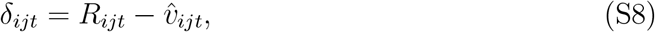

which is the difference between the actual perceived reward *R*_*ijt*_ and the estimated value 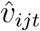 The learning updates for the *θ* parameters are given by

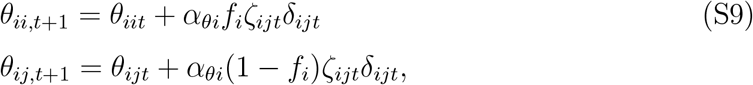

where

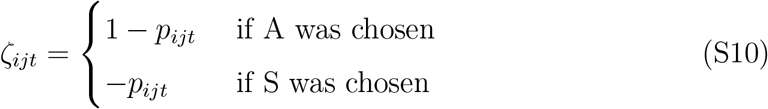

is referred to as a policy-gradient factor and *α*_*θi*_ is the preference learning rate for individual *i*. Note that *ζ*_*ijt*_ will be small if *p*_*ijt*_ is close to one and individual *i* performed action A, which slows down learning, with a corresponding slowing down if *p*_*ijt*_ is close to zero and S is chosen. There are also learning updates for the *w* parameters given by

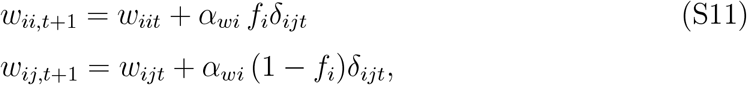

where *α*_*wi*_ is the value learning rate for individual *i*.

The updates to the policy parameters *θ* can be described using derivatives of the logarithm of the probability of choosing an action with respect to the parameters. Using equation (S2), we obtain

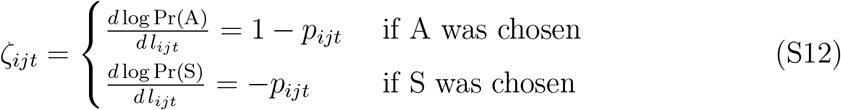

for the derivative of the logarithm of the probability of choosing an action, A or S, with respect to the preference for A, which corresponds to equation (S10). From equation (S3) it follows that

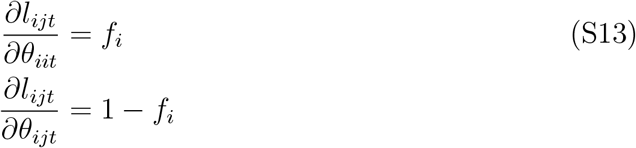

and this gives the learning updates of the *θ* parameters in equation (S9). The updates of the *w* parameters of the value function can also be described using derivatives. From equation (S4) it follows that

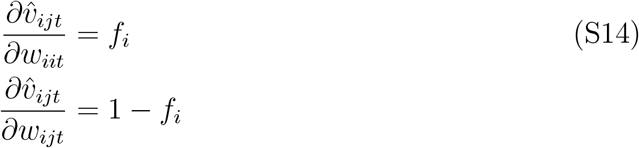

and this gives the learning updates of the *w* parameters in equation (S11).

### Bystander updates

As in previous models [8, 9], bystander effects are modelled as observational learning. When there is a dominance interaction in a group, individuals other then the interacting pair *i* and *j*, for instance an individual *k*, can use the outcome to update the learning parameters. Assume that individual *k* only performs this updating if *i* and *j* end their interaction by using AS or SA (because there is no clear ‘winner’ in AA and SS interactions, and bystanders do not perceive the costs of AA interactions). The probabilities for individuals *i* and *j* to use A are *p*_*ijt*_ and *p*_*jit*_, from equation (S2). These are ‘true’ values and are not known by individual *k*. However, given that the outcome is either AS or SA, one readily derives that the logit of the probability that it is AS is

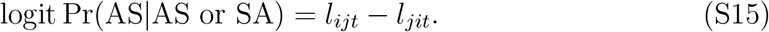

From equation (S3) one can see that this involves various learning parameters for *i* and *j*. For bystander learning an assumption is needed about how an individual *k* represents this logit. A simple assumption is that *k* represents the logit as

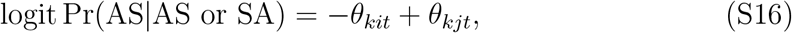

which entails that *k* does not use any information about *q*_*i*_ or *q*_*j*_. The assumption is reasonable in that a large *θ*_*kit*_ means that *k* behaves as if individual *i* is weak, and similarly for *θ*_*kjt*_. Using the notation

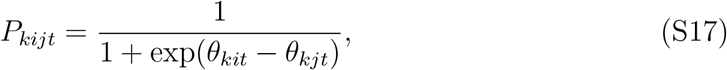

one possibility for the bystander updates by *k* is

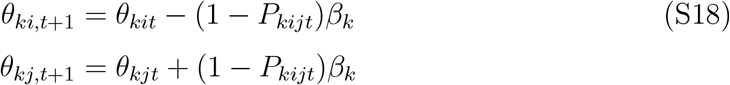

if *i* wins (outcome is AS) and

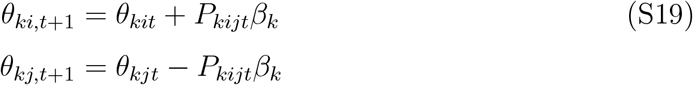

if *j* wins (outcome is SA). The parameter *β*_*k*_ is a measure of how salient or significant a bystander observation is for individual *k*, and this parameter is assumed to be genetically determined and can evolve. These bystander updates are similar to the direct-learning updates of the actor component of the actor-critic model and were used previously [8].

There is also the possibility that the salience for a bystander of a contest outcome is influenced by additional information the bystander might have, either from current or from previous observations. For instance, if the observations in equation (S1) are about relative size, a bystander might have estimates *ξ*_*ki*_ and *ξ*_*kj*_ of its own size in relation to the contestants *i* and *j*. Instead of the bystander updates above we might then have

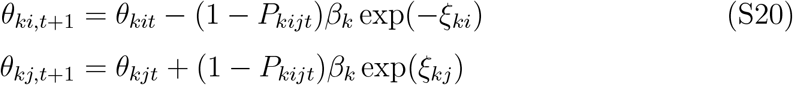

if *i* wins (outcome is AS) and

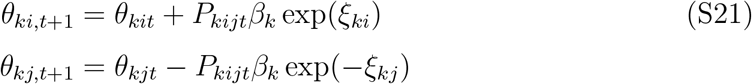

if *j* wins (outcome is SA). These updates entail that a bystander *k* pays particular attention to wins by an individual perceived to be bigger than itself, and to losses by an individual perceived to be smaller. These updates were used previously [9] and we also use them in our simulations here.

### Contests

If a dominance relation has already been established between contestants *i* and *j*, there is no interaction. If not, the contestants go through a number of rounds, at minimum 10 rounds and at maximum 200 rounds of interaction. If there are 5 successive rounds where *i* uses A and *j* uses S (5 AS rounds), the contest ends and *i* is considered dominant over *j*, and vice versa if there are 5 successive SA rounds. Further, the contest ends in a draw if there are 5 successive SS rounds.

### Details of dominance ranking

The ranking is among surviving individuals and is based of how many other group members an individual dominates (this measure has been referred to as a score structure [47]). If some individuals dominate the same number of other group members, their relative rank is randomly determined (this happened occasionally in our simulations). As an extreme example, if all individuals would use action S in the contests, there would be no real dominance hierarchy, because each would dominate zero other group members, and all ranks would be randomly determined (this never happened in our simulations).

### Acquired resources and interference as a function of rank

For the competing sex, a surviving group member of rank *k* obtains an amount *V* (*k*) of contested AR, which is further modified by interference by dominants. Our assumptions of the amounts without interference can be seen from the shapes of the curves labelled *L*_1_ and *G*_1_ in Fig. 1c. An individual *i* with interference trait *κ*_*i*_ pays a cost of performing interference by multiplying its contested AR by a fraction

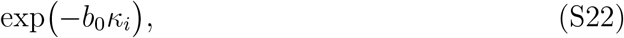

provided the individual has others that are subordinate to it. This is illustrated by the effect on self curve in Fig. 1b. Each of the subordinates exposed to interference has its contested AR changed through multiplying by a fraction

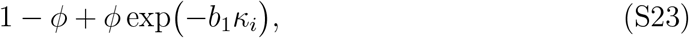

which is illustrated by the effect on others in Fig. 1b. An individual that is subordinate to more than one dominant suffers from interference from each dominant. We assume that the effects of interference on contested AR interact multiplicatively. The overall effects of this for different values of the strength of interference are shown in Fig. 1b. Our model assumptions represent just one possibility, and would likely need to be modified to describe a particular case of interference in nature.

### Mortality and reduced parenting ability from fighting damage

An individual with accumulated damage *D*_*it*_ survives from contests to reproduction with probability

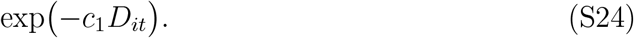

The parenting ability of a surviving individual is

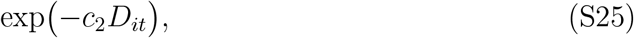

and is illustrated in Fig. 1d. For contested RS, the interpretation is that reduced parenting ability corresponds to an effective reduction of AR. For contributions to outside-option reproduction, reduced parenting ability correspond to a reduction in the probability of contributing.

## References

[1] Paula Stockley and Jakob Bro-Jørgensen. Female competition and its evolution-ary consequences in mammals. Biological Reviews, 86(2):341–366, 2011. doi: 10.1111/j.1469-185X.2010.00149.x.

[2] T. Clutton-Brock and E. Huchard. Social competition and its consequences in female mammals. Journal of Zoology, 289(3):151–171, 2013. doi: 10.1111/jzo.12023.

[3] T. H. Clutton-Brock and E. Huchard. Social competition and selection in males and females. Philosophical Transactions of the Royal Society B: Biological Sciences, 368(1631):20130074, 2013. doi: 10.1098/rstb.2013.0074.

[4] Lee Ellis. Dominance and reproductive success among nonhuman animals: A cross-species comparison. Ethology and Sociobiology, 16(4):257–333, 1995. doi: 10.1016/0162-3095(95)00050-U.

[5] T. H Clutton-Brock. Reproductive skew, concessions and limited control. Trends in Ecology & Evolution, 13(7):288–292, 1998. doi: 10.1016/S0169-5347(98)01402-5.

[6] Ilik Saccheri and Ilkka Hanski. Natural selection and population dynamics. Trends in Ecology & Evolution, 21(6):341–347, 2006. doi: 10.1016/j.tree.2006.03.018.

[7] John M. McNamara and Olof Leimar. Game Theory in Biology: Concepts and Frontiers. Oxford University Press, Oxford (in press), 2020.

[8] Olof Leimar. The evolution of social dominance through reinforcement learning. The American Naturalist, 197(5):560–575, 2021. doi: 10.1086/713758.

[9] Olof Leimar and Redouan Bshary. The influence of reproductive skew on fighting costs and winner-loser effects in social dominance evolution. Journal of Animal Ecology, 2022. doi: 10.1111.1365-2656.13691.

[10] John Maynard Smith and G. R. Price. The logic of animal conflict. Nature, 246 (5427):15–18, 1973. doi: 10.1038/246015a0.

[11] Christian S. Willisch and Peter Neuhaus. Alternative mating tactics and their impact on survival in adult male alpine ibex (Capra ibex ibex). Journal of Mammalogy, 90(6):1421–1430, 2009. doi: 10.1644/08-MAMM-A-316R1.1.

[12] Christian S. Willisch and Peter Neuhaus. Social dominance and conflict reduction in rutting male Alpine ibex, Capra ibex. Behavioral Ecology, 21(2):372–380, 2010. doi: 10.1093/beheco/arp200.

[13] Sam K. Patterson, Shirley C. Strum, and Joan B. Silk. Resource competition shapes female–female aggression in olive baboons, Papio anubis. Animal Behaviour, 176:23–41, 2021. doi: 10.1016/j.anbehav.2021.03.013.

[14] Ted J. Case and Michael E. Gilpin. Interference competition and niche theory. Proceedings of the National Academy of Sciences, 71(8):3073–3077, 1974. doi: 10.1073/pnas.71.8.3073.

[15] Vincent Le Bourlot, Thomas Tully, and David Claessen. Interference versus exploitative competition in the regulation of size-structured populations. The American Naturalist, 184(5):609–623, 2014. doi: 10.1086/678083.

[16] Wouter K. Vahl, Tamar Lok, Jaap van der Meer, Theunis Piersma, and Franz J. Weissing. Spatial clumping of food and social dominance affect interference competition among ruddy turnstones. Behavioral Ecology, 16(5):834–844, 2005. doi: 10.1093/beheco/ari067.

[17] T. H. Clutton-Brock and G. A. Parker. Punishment in animal societies. Nature, 373(6511):209–216, 1995. doi: 10.1038/373209a0.

[18] Stuart A. West, Andy Gardner, David M. Shuker, Tracy Reynolds, Max Burton-Chellow, Edward M. Sykes, Meghan A. Guinnee, and Ashleigh S. Griffin. Cooperation and the scale of competition in humans. Current Biology, 16(11):1103–1106, 2006. doi: 10.1016/j.cub.2006.03.069.

[19] Nichola J. Raihani and Redouan Bshary. Punishment: one tool, many uses. Evolutionary Human Sciences, 1:e12, 2019. doi: 10.1017/ehs.2019.12.

[20] A. Gardner and S. A. West. Spite and the scale of competition. Journal of Evolutionary Biology, 17(6):1195–1203, 2004. doi: 10.1111/j.1420-9101.2004.00775.x.

[21] Stuart A. West and Andy Gardner. Altruism, spite, and greenbeards. Science, 327(5971):1341–1344, 2010. doi: 10.1126/science.1178332.

[22] Julia Ostner, Charles L. Nunn, and Oliver Schülke. Female reproductive synchrony predicts skewed paternity across primates. Behavioral Ecology, 19(6):1150–1158, 2008. doi: 10.1093/beheco/arn093.

[23] Dieter Lukas and Elise Huchard. The evolution of infanticide by females in mam-mals. Philosophical Transactions of the Royal Society B: Biological Sciences, 374 (1780):20180075, 2019. doi: 10.1098/rstb.2018.0075.

[24] Yayong Wu, Martin J. Whiting, Jinzhong Fu, and Yin Qi. The driving forces behind female-female aggression and its fitness consequence in an Asian agamid lizard. Behavioral Ecology and Sociobiology, 73(6):73, 2019. doi: 10.1007/s00265-019-2686-8.

[25] Alain Houle and Richard W. Wrangham. Contest competition for fruit and space among wild chimpanzees in relation to the vertical stratification of metabolizable energy. Animal Behaviour, 175:231–246, 2021. doi: 10.1016/j.anbehav.2021.03.003.

[26] H Caswell. Matrix population models, second edition. Sinauer Associates, Inc., Sunderland, MA, 2001.

[27] Bruce Wallace. Hard and soft selection revisited. Evolution, 29(3):465–473, 1975. doi: 10.1111/j.1558-5646.1975.tb00836.x.

[28] Jens Jacob, Nur Aini Herawati, Stephen A. Davis, and Grant R. Singleton. The impact of sterilized females on enclosed populations of ricefield rats. Journal of Wildlife Management, 68(4):1130–1137, 2004. doi: 10.2193/0022-541X(2004)068[1130:TIOSFO]2.0.CO;2.

[29] Jens Jacob, Grant R. Singleton, Lyn A. Hinds, Jens Jacob, Grant R. Singleton, and Lyn A. Hinds. Fertility control of rodent pests. Wildlife Research, 35(6): 487–493, 2008. doi: 10.1071/WR07129.

[30] Layne G. Adams, Robert O. Stephenson, Bruce W. Dale, Robert T. Ahgook, and Dominic J. Demma. Population dynamics and harvest characteristics of wolves in the central brooks range, alaska. Wildlife Monographs, 170(1):1–25, 2008. doi: 10.2193/2008-012.

[31] Cristina Sartori, Serena Mazza, Nadia Guzzo, and Roberto Mantovani. Evolution of increased competitiveness in cows trades off with reduced milk yield, fertility and more masculine morphology. Evolution, 69(8):2235–2245, 2015. doi: 10.1111/evo.12723.

[32] Marta C. Soares, Renata Mazzei, Sónia C. Cardoso, Cândida Ramos, and Redouan Bshary. Testosterone causes pleiotropic effects on cleanerfish behaviour. Scientific Reports, 9(1):15829, 2019. doi: 10.1038/s41598-019-51960-w.

[33] IM Côte. Evolution and ecology of cleaning symbioses in the sea. In RN Gibson and M Barnes, editors, Oceanography and Marine Biology, volume 38, pages 311–355. 2000.

[34] D. R. Robertson. Social control of sex reversal in a coral-reef fish. Science, 177 (4053):1007–1009, 1972. doi: 10.1126/science.177.4053.1007.

[35] Noah Snyder-Mackler, Joseph Robert Burger, Lauren Gaydosh, Daniel W. Belsky, Grace A. Noppert, Fernando A. Campos, Alessandro Bartolomucci, Yang Claire Yang, Allison E. Aiello, Angela O’Rand, Kathleen Mullan Harris, Carol A. Shively, Susan C. Alberts, and Jenny Tung. Social determinants of health and survival in humans and other animals. Science, 368(6493):eaax9553, 2020. doi: 10.1126/science.aax9553.

[36] Martin N. Muller, Drew K. Enigk, Stephanie A. Fox, Jordan Lucore, Zarin P. Machanda, Richard W. Wrangham, and Melissa Emery Thompson. Aggression, glucocorticoids, and the chronic costs of status competition for wild male chimpanzees. Hormones and Behavior, 130:104965, 2021. doi: 10.1016/j.yhbeh.2021.104965.

[37] R. A. Johnstone and M. A. Cant. Reproductive skew and the threat of eviction: a new perspective. Proceedings of the Royal Society B: Biological Sciences, 266 (1416):275–279, 1999. doi: 10.1098/rspb.1999.0633.

[38] Rufus A. Johnstone. Models of reproductive skew: A review and synthesis. Ethology, 106(1):5–26, 2000. doi: https://doi.org/10.1046/j.1439-0310.2000.00529.x.

[39] Peter Nonacs. Tug-of-war has no borders: it is the missing model in reproductive skew theory. Evolution, 61(5):1244–1250, 2007. doi: 10.1111/j.1558-5646.2007.00092.x.

[40] Markus Port and Peter M. Kappeler. The utility of reproductive skew models in the study of male primates, a critical evaluation. Evolutionary Anthropology, 19 (2):46–56, 2010. doi: https://doi.org/10.1002/evan.20243.

[41] Peter Nonacs and Reinmar Hager. The past, present and future of reproductive skew theory and experiments. Biological Reviews, 86(2):271–298, 2011. doi: 10.1111/j.1469-185X.2010.00144.x.

[42] Michael A. Cant and Andrew J. Young. Resolving social conflict among females without overt aggression. Philosophical Transactions of the Royal Society B: Biological Sciences, 368(1631):20130076, 2013. doi: 10.1098/rstb.2013.0076.

[43] John Maynard Smith. On Evolution. Edinburgh University Press, Edinburgh, 1972.

[44] J. Maynard Smith. Evolution and the Theory of Games. Cambridge University Press, Cambridge, 1982.

[45] G. A. Parker. Assessment strategy and the evolution of fighting behaviour. Journal of Theoretical Biology, 47(1):223–243, 1974. doi: 10.1016/0022-5193(74)90111-8.

[46] Richard S Sutton and Andrew G Barto. Reinforcement learning: An introduction second edition. MIT Press, Cambridge, MA, 2018.

[47] H. G. Landau. On dominance relations and the structure of animal societies: I. Effect of inherent characteristics. The Bulletin of Mathematical Biophysics, 13(1): 1–19, 1951. doi: 10.1007/BF02478336.

